# SARS-CoV-2 Spike Pseudoviruses: A Useful tool to study virus entry and address emerging neutralization escape phenotypes

**DOI:** 10.1101/2021.07.16.452709

**Authors:** Raj Kalkeri, Zhaohui Cai, Shuling Lin, John Farmer, Yury V. Kuzmichev, Fusataka Koide

**Affiliations:** Department of Infectious Disease Research, Drug Development Division, Southern Research, Frederick, MD 21701, USA;, (R.K.); (Z.C.); (S.L); (Y.V.K.); (F.K.); Department of Infectious Disease Research, Drug Development Division, Southern Research, Birmingham, AL 35205, USA; (J.F.)

**Keywords:** COVID-19, SARS-CoV-2, pseudovirus, Variants of Concern, neutralizing antibodies

## Abstract

SARS-CoV-2 genetic variants are emerging around the globe. Unfortunately, several SARS-CoV-2 variants, especially, variants of concern (VOC) are less susceptible to neutralization by the convalescent and post-vaccination sera, raising concerns of increased disease transmissibility and severity. Recent data suggests the SARS-CoV-2 neutralizing anti-body levels as a good correlate of vaccine mediated protection. However, currently used BSL3 based virus micro-neutralization (MN) assays are more laborious, time consuming and expensive, underscoring the need for BSL2 based, cost effective neutralization assays against SARS-CoV-2 variants. In light of this unmet need, we have developed a BSL2 pseudovirus based neutralization assay (PBNA) in cells expressing Angiotensin Converting Enzyme-2 (ACE2) receptor for SARS-CoV-2. The assay is reproducible (R^2^=0.96), demonstrates a good dynamic range and high sensitivity. Our data suggests that the biological Anti-SARS-CoV-2 research reagents such as NIBSC 20/130 show lower neutralization against B.1.351 RSA and B1.1.7 UK VOC, whereas a commercially available monoclonal antibody MM43 retains activity against both these variants. SARS-CoV-2 Spike Pseudovirus based neutralization assays for VOC would be useful tools to measure the neutralization ability of candidate vaccines in both preclinical models and clinical trials and further help develop effective prophylactic countermeasures against emerging neutralization escape phenotypes.

## 1. Introduction

Severe acute respiratory syndrome coronavirus 2 (SARS-CoV-2), the causative agent of coronavirus dis-ease 2019 (COVID-19), remains a major global health challenge responsible for more than 4-million deaths since the start of the pandemic in December 2019, with an estimated 187 million confirmed cases as of 12 July 2021 [1]. While the development of novel prophylactic and therapeutic measures, such as vaccines, monoclonal antibodies, and antiviral medications have been instrumental in slowing down the pace of the pandemic, emergence of several variants of concern (VOC) has raised issues about potential immune escape [2]. SARS-CoV-2, one of the largest known RNA viruses with ^~^29.8 kb genome, is an enveloped, positive-sense single-stranded RNA virus, belonging to the genus of β-coronaviruses of the coronaviridae family [3]. The spike (S) glycoprotein, present on the surface of the virion is responsible for host cell attachment and fusion with the human angiotensin-converting enzyme 2 (hACE2) receptor. S glycoprotein is currently considered a major immunogenic component of SARS-CoV-2 [4–6]. Measuring the effect of neutralizing antibodies, an important correlate of protection, against the S glycoprotein is of primary importance in fighting the pandemic. Currently a number of clinical trials investigating such therapeutic interventions are ongoing [7].

In order to determine the efficacy of approaches for measuring the effect of neutralization against emerging variants of concern, assays that are capable of measuring serological responses to the spike glycoprotein are of extreme importance. Current assays used for such purposes rely on principles of microneutralization (MN) or enzyme-linked immunosorbent assays (ELISA), and several ELISA derivatives [8–10]. SARS-CoV-2 MN assays relying on the neutralization of a wild-type, replicating virus, are considered the gold-standard methods for the evaluation of neutralization ability of coronavirus induced antibodies. However, in its present form, MN assay is expensive and labor intensive, requiring the use of a biosafety level three containment (BSL-3), making it challenging to adapt into large scale clinical trials [11]. At the other extreme are ELISA methods. While considered safe and adaptable to high-throughput format, however, they measure total antibodies against the protein and do not measure the neutralization titers in contrast to the MN formats [9,12–14].

To avoid the use of highly restrictive BSL-3 environment and improve upon the ELISA, the implementation of replication-deficient pseudo-viruses containing the viral coat proteins of interest has been suggested as a safe and useful alternative [15]. Pseudovirus-based platforms have been successfully employed in the study of highly infectious and pathogenic viruses, including Ebola, Middle Easter Respiratory Syndrome (MERS), rabies, Marburg, Lassa, and others [16–20]. Recently, a number of groups have successfully generated SARS-CoV-2 pseudo-viruses using murine leukemia virus (MLV), vesicular stomatitis virus (VSV), as well as human immunodeficiency virus (HIV) platforms and employed them for the evaluation of neutralizing antibodies through different readout systems [21–27]. However, there is currently limited data on the use of such systems in the evaluation of the SARS-CoV-2 variants of concern. To address this, we developed and optimized a robust pseudovirus-based neutralization assay (PBNA) and evaluated it against SARS-CoV-2 614D and two variants of concern (B1.1.7, UK variant, B1.351 RSA variant). Furthermore, we evaluated the performance characteristics of the positive plasma control (NIBSC 20/130) and two commercially available monoclonal antibodies in PBNA. Here, we present detailed methods and the performance of PBNA, which could be adapted for use in various quantitative, medium/high-throughput virus neutralization screens in a standard BSL-2 laboratory environment.

## 2. Materials and Methods

### 2.1. Pseudoviruses

The virus backbone (HIV-1 NL4-3 ΔEnv Vpr Luciferase Reporter Vector, pNL4-3.Luc.R-E-, NIH AIDS Reagent, Catalog Number: 3418 was licensed/obtained from the New York University School of Medicine. Codon Optimized SARS-CoV-2 Spike gene from isolate 2019-nCoV_HKU-SZ-002a_2020 (GenBank: MN938384) was synthesized at GeneWiz, South Plainfield, NJ. Spike gene was cloned into eukaryotic expression plasmid pcDNA3.1 to generate plasmid, designated as pSRC332. For pseudovirus production, Lenti-X-293T cells were co-transfected with pNL4-3.Luc.R-E- and pSRC322 using JetPrime transfection reagent (Polyplus Transfection, New York, NY). Briefly, 6×106 Lenti-X-293T cells were seeded in T75 flask one day before transfection. The next day cells were co-transfected with 3 μg pSRC332 and 12 μg pNL43. Luc. R. E using JetPrime^®^ Transfection Reagent following the manufacturer’s instruction. Five hours later, plasmid DNA-transfection complexes were removed and cells were washed with PBS once and then new culture medium was added. Seventy-two hours post transfection, SARS-CoV-2 pseudoviruses containing culture supernatant was harvested and clarified by centrifugation at 1200 RPM for 5minutes and stored at −70°C in 0.5mL aliquots until use. A set of murine lentivirus (MLV) based pseudo-viruses (replication-deficient MLV) for the 614D (Cat#MBS434275), B1.357 RSA variant (Cat#MBS434287) and B1.1.7 UK Variant (Cat# MBS434286) were purchased from MyBioSource Inc., San Diego, USA.

### 2.2. Serum and Monoclonal antibodies

NIBSC 20/130 (National Institute of Biological Standards, Ridge, UK), Anti-SARS-CoV-2 RBD Neutralizing Antibody, Human IgG1 (Cat#SAD-S35, Acro Biosystems, Newark, DE), SARS-CoV-2 (2019-nCoV) spike neutralizing mouse monoclonal antibody (Cat#40591-MM43, Sino Biological, Wayne, PA) were used as positive controls in the assay. Sera from unvaccinated healthy monkeys were used as negative controls. Sera 28-days post-infection from three (3) different species of non-human primates (NHPs) rhesus macaques (RM), cynomolgus monkeys (CM), African Green Monkeys (AGMs) (N=2 each), challenged with SARS-CoV-2 (SARS-CoV-2 USA_WA1/2020 strain at 4.0 x 10^6^ TCID_50_/mL through intra-tracheal route) were used in the pseudovirus assay.

### 2.3. Pseudovirus Based Neutralization Assay

HEK293T-ACE2 cells (NR-52511, BEI Resources, Manasas, VA) were grown in Dulbecco’s Minimal Essential Medium (DMEM, Lonza, Walkersville, MD), supplemented with 10% Fetal Bovine Serum (FBS) according to standard culture conditions. Cells were subcultured twice a week at a split ratio of 1:5 to 1:10 using standard cell culture techniques and the cell culture media as specified below in Table 2. Total cell number and percent viability determinations were performed using a hemocytometer and trypan blue dye exclusion. Cell viability was greater than 95% for the cells to be utilized in the assays. HEK293-ACE2 cells were seeded at ^~^20,000 cells per well into 96-well plates (100 μL per well) DMEM with 10% FBS without antibiotic the day before the assay. Next day, test serum heat inactivated at 56°C for 30 minutes was serially diluted 5-fold in DMEM+2% FBS, followed by mixing with equal volume of SARS-CoV-2 pseudoviral particles (final volume of 100 μL) and incubation at 37°C for 1 hour. Media from the cell culture was removed, 100 μL of serum-virus mixture was added into each well. Each assay plate contained cells without pseudovirus infection (Cell control/background) and pseudovirus only control (Virus Control). Each assay run also included a positive control (serum or antibody with known neutralizing activity). Cell culture plate was centrifuged at 700 rpm for 15 min at 4°C, followed by incubation for 72 hrs, at 37 °C with 5% CO_2_. Luciferase activity in the infected cells was measured by removing the culture supernatant and adding 50 μL Luciferase assay reagent (Firefly luciferase reagent, Promega. Madison, WI). Luminescence was recorded using the luminescence plate reader (Clariostar plate reader, BMG LabTech Inc, Cary, NC). Luminescence signal was normalized to the virus control after background subtraction and analyzed using PRISM software using four parameter logistic curve fitting to calculate 50% pseudovirus based neutralization index (PBNI_50_).

## 3. Results

Pseudovirus based neutralization assay (PBNA) for SARS-CoV-2 spike protein variants (614D, B1.1.7 and B1.357) was developed in HEK293T-cells expressing ACE2 receptor for SARS-CoV-2. A set of positive controls (NIBSC 20/130, Acro-SAD-S35 and Sino-40591-MM43) were tested in this assay system. We demonstrate that NIBSC 20/130 shows significantly reduced neutralization against B1.357 RSA and B1.1.7 UK variants, whereas Sino-40591-MM43 show better neutralization against these variants.

### 3.1. Dose dependence of SARS-CoV-2 spike protein pseudovirus variants

To develop the pseudovirus based assay, HEK293T-hACE2 cells were infected with increasing concentrations of SARS-CoV-2 spike protein pseudovirus variants. Luciferase signal in the infected cells was analyzed after 3-days post-infection. As shown in Figure 1, there was a dose dependent increase in signal for all three pseudo-viruses with a dynamic range of 2-logs. Pseudoviruses for 614D and B1.1.7 UK variant showed similar luciferase signal, (signal of 188,869.5±13,500.8 and 66,315±1,640.5 RLU) at the highest concentration tested (40 μL/well). This resulted in a robust signal to background (S/B) ratio in the range of 2337 to 6656. In contrast, cells infected with B1.351 RSA variant showed significantly lower signal (9400.5 RLU, 1-log lower) compared to the 614D and B1.1.7 UK variant pseudoviruses, at the highest concentration tested. Based upon these results and the optimal S/B level, we chose the virus inoculum for 614D and B1.1.7 UK and B1.357 RSA variant for testing the serum samples to detect neutralizing antibodies.

**Figure 1.**
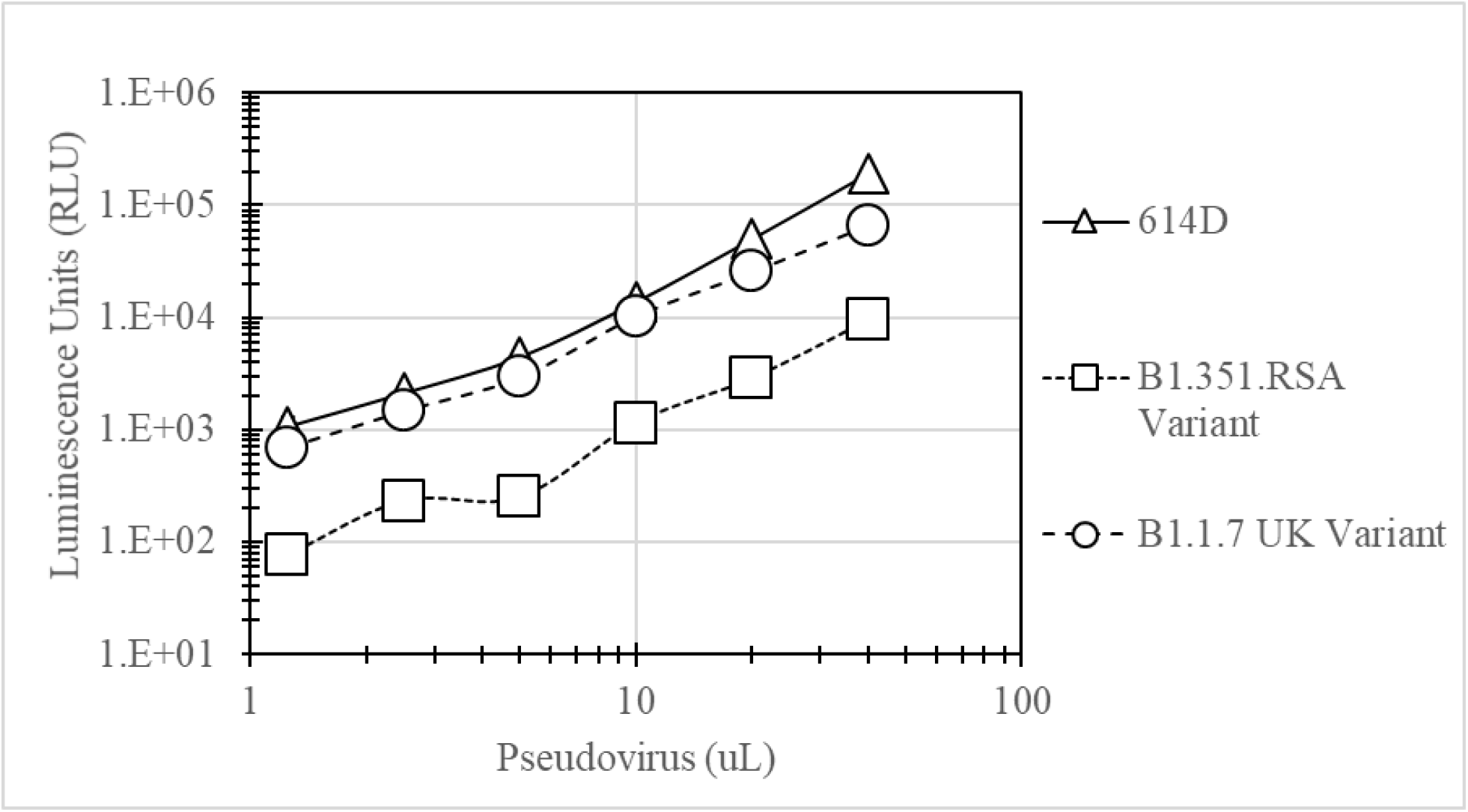
Dose dependence of SARS-CoV-2 spike protein pseudo-viruses: HEK293T-hACE2 cells seeded in 96 well plates were infected with increasing concentrations of SARS-CoV-2 spike protein pseudo-viruses. Intracellular luciferase signal (as a marker of spike protein mediated pseudovirus infection) measured at the end of 3 days is shown on the Y-axis. Triangles-614D, Circles-B1.1.7 UK variant, Squares-B1.351 RSA Variant.

### 3.2. Suitability of PBNA for evaluation of neutralization antibodies and Reproducibility of the SARS-CoV-2 spike protein pseudovirus assay

It is important for the assays to be reproducible between different experiments to consistently determine the neutralizing antibodies in the test serum samples. To address this question, we tested 6 serum samples collected 28-days post-infection from different species of monkeys (Rhesus Macaques, Cynomolgus monkeys, African Green Monkeys, N=2 each) challenged with SARS-CoV-2 in the pseudovirus based neutralization assay (PBNA) as mentioned in the methods section. Serum samples from all three monkey species showed PBNI_50_ ranging from 147 to 732.7. Human IgG1Anti-SARS-CoV-2 Spike RBD Neutralizing Antibody (Acro-SAD-S35) and a mouse monoclonal antibody SARS-CoV-2 (2019-nCoV) Spike Neutralizing Antibody (Sino-40591-MM43) were used as positive controls in the assay. A non-challenged monkey serum was used as a negative control in the assay. The positive controls showed robust inhibition of the pseudovirus infection in PBNA with 50% pseudovirus based neutralization index (PBNI_50_) of 11,164.0 and 15,706.0 respectively. Negative control showed a PBNI_50_ of 123.3 ± 54.2 in the assay, which might be due to preexisting antibodies (may be due to prior exposure to other Coronaviruses). More than 2-log difference between the negative and positive controls confirm the significant assay dynamic range, which could enable screening antibodies with ability to differentiate between samples with varying neutralization potencies against SARS-CoV-2.

The assay was repeated on two different days by the same operator. Data from both the experiments were compared and shown in Figure 2. Positive controls showed robust activity in both the assays with mean ± SD of 10,522.5 ± 907.2 and 14,944 ± 1,077.6 in the assay. Both sets of data showed good correlation with an R^2^ value of 0.96, suggesting that the assay is reproducible. The data also suggests that all three monkey species (Rhesus Macaques, Cynomolgus monkeys, African Green Monkeys) developed neutralizing antibodies (Mean ± SD of 298.7 ± 228.9, 313.9 ± 98.8, 665.1 ± 191.7) respectively.

**Figure 2.**
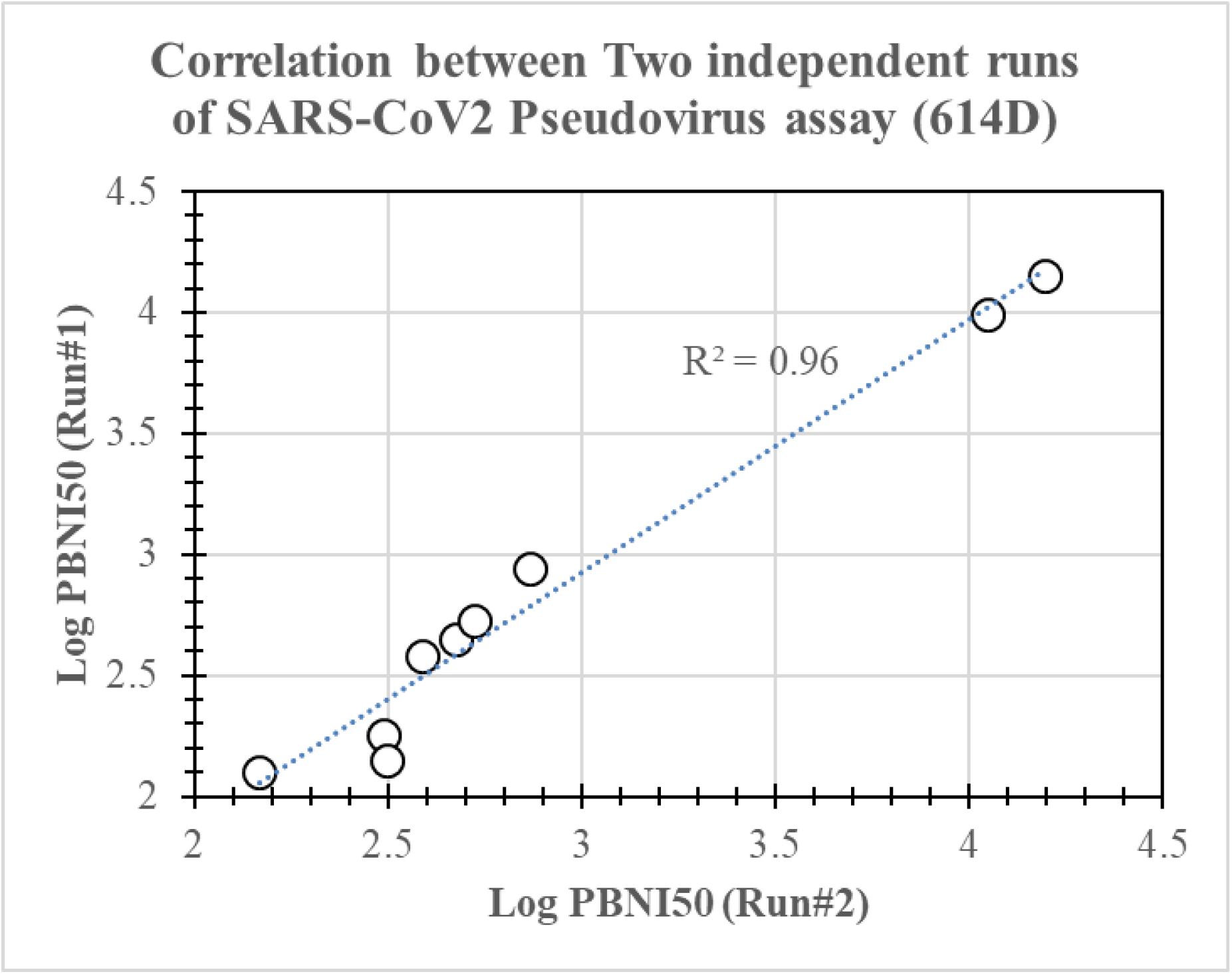
Reproducibility of the SARS-CoV-2 spike protein pseudovirus based neutralization assay (PBNA): PBNA for 6-serum samples from SARS-CoV-2 challenged NHP, 2 positive and 1-unchallenged NHP controls was conducted in two independent experiments (11th and 18th June2021). Log transformed PBNI_50_ from each experiment is plotted on X- and Y-axis respectively. Correlation of R^2^=0.96, with the trend line between the PBNI_50_ from both the experiments is shown by the dotted line in the figure.

### 3.3. SARS-CoV-2 challenged NHP serum samples show neutralization against SARS-CoV-2 614D and B1.1.7 UK Variants

Recently SARS-CoV-2 variants of concerns are being reported in the literature. There are some concerns about the differential neutralizing abilities induced by SARS-CoV-2 infection against variants of concern. To evaluate the neutralizing antibodies against the variants of concern, we tested 6-serum samples from monkeys challenged with SARS-CoV-2 USA_WA1/2020 strain against pseudoviruses expressing spike proteins of B1.315 RSA and B1.1.7 UK variants, in PBNA. As shown in Figure 3, these serum samples showed similar neutralizing antibodies against both 614D and B1.1.7 UK variant. For B1.1.7 UK variant, PBNI_50_ ranged from 49.6 to 811.5. Interestingly, majority of the serum samples (5 out of 6) showed lower amounts of neutralizing antibodies (PBNI_50_ <100) against B1.315 RSA. Only one animal (3F16765) showed PBNI_50_ of 449.2 against B1.315 RSA, which was similar to both 614D and B1.1.7 UK variant PBNI_50_ levels. These data suggest that neutralizing antibodies elicited by SARS-CoV-2 USA_WA1/2020 strain are effective against B1.1.7 UK variant, but less effective against B1.315 RSA variant.

**Figure 3.**
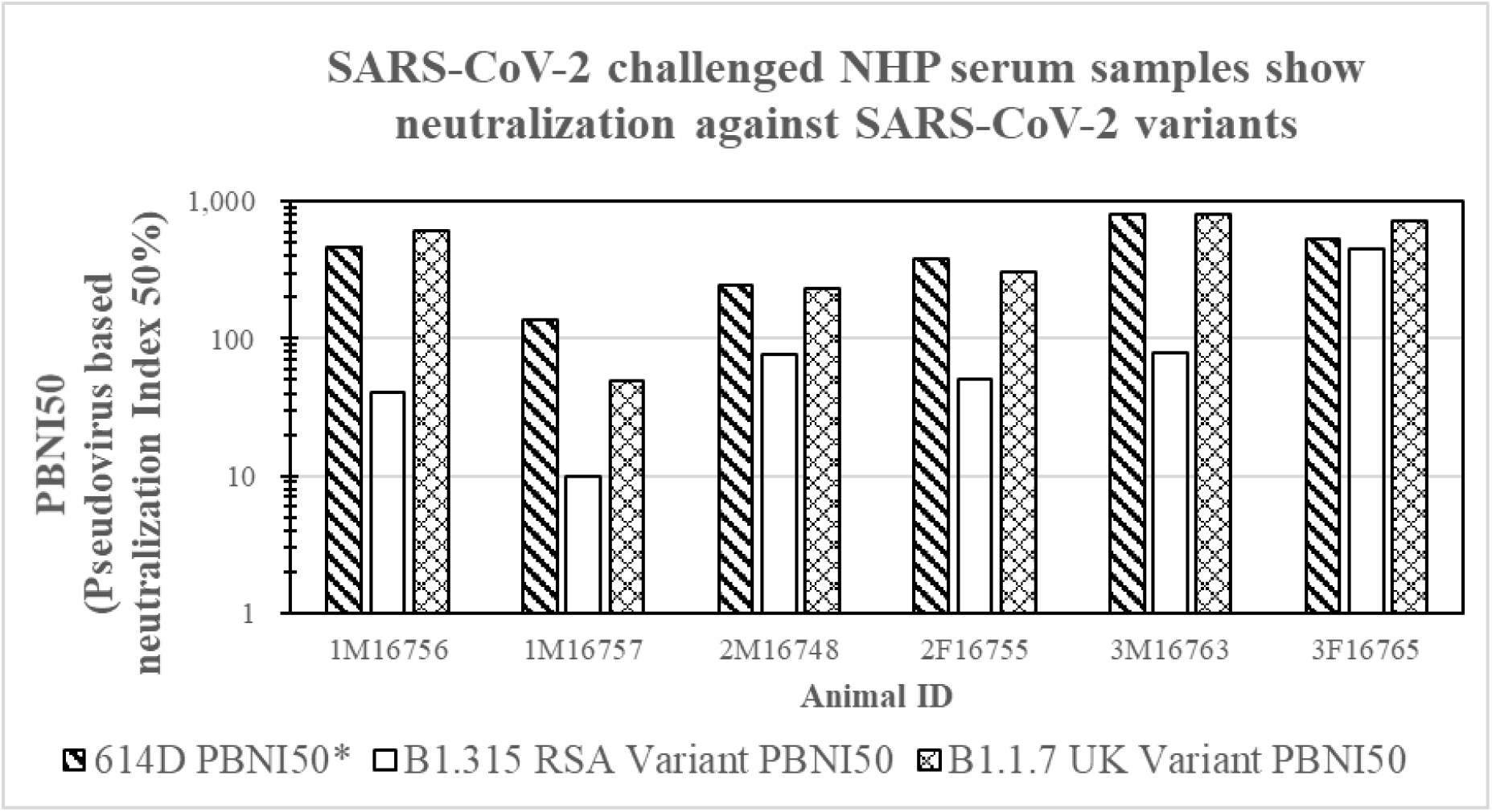
SARS-CoV-2 challenged NHP serum samples show neutralization against SARS-CoV-2 Variants: Six (6)-serum samples from SARS-CoV-2 challenged NHPs [N=2 from each species: Rhesus Macaques (1M16756, 1M16757), Cynomolgus monkeys (2M16748, 2F16748), African green monkeys (3M16763, 3F16765)] were subjected to PBNA using the pseudoviruses for 614D, and variants as described in the methods section. PBNI_50_ for each animal is shown on the Y-axis. Cross hatch filled bars show average PBNI_50_ from two independent experiments for 614D, plain bars show PBNI_50_ for B1.315 RSA variant, diamond filled bars show PBNI_50_ for B1.1.7 UK Variant.

### 3.4. Linearity of neutralization index (PBNI50) generated by the pseudovirus assay

In order to identify the linearity of neutralization index (PBNI_50_), three NHP serum samples with high PBNI_50_ (from animals 1M16756, 3M16763, 3F16765), were serially diluted (5-fold) to generate test samples with high, medium and low PBNI_50_. A positive control (Acro-SAD-S35) similarly diluted was also used as a positive control in the assay. Nine (9)-test samples (3 each of high, medium and low PBNI_50_ samples), were tested in PBNA. Results shown in Figure. 4 reveal a linear reduction in the PBNI_50_ titers. High, medium and low PBNI_50_ test groups showed PBNI_50_ (Mean ± SD) of 504.7 ± 156.3, 108.6 ± 20.2 29.5 ± 16.2 respectively. Slope analysis revealed a robust R^2^ value of 0.9978 suggesting a linear relationship between both expected and measured PBNI_50_ values. Data analysis of the positive control also showed a linear range of PBNI_50_ for high, medium and low PBNI_50_ samples ranging from 6819 to 115.5 with R^2^ value of 0.989 (data not shown). Besides linearity of dilution, this data also revealed the sensitivity of the PBNA with lower PBNI_50_ detection of 29.5 ± 16.2.

**Figure 4.**
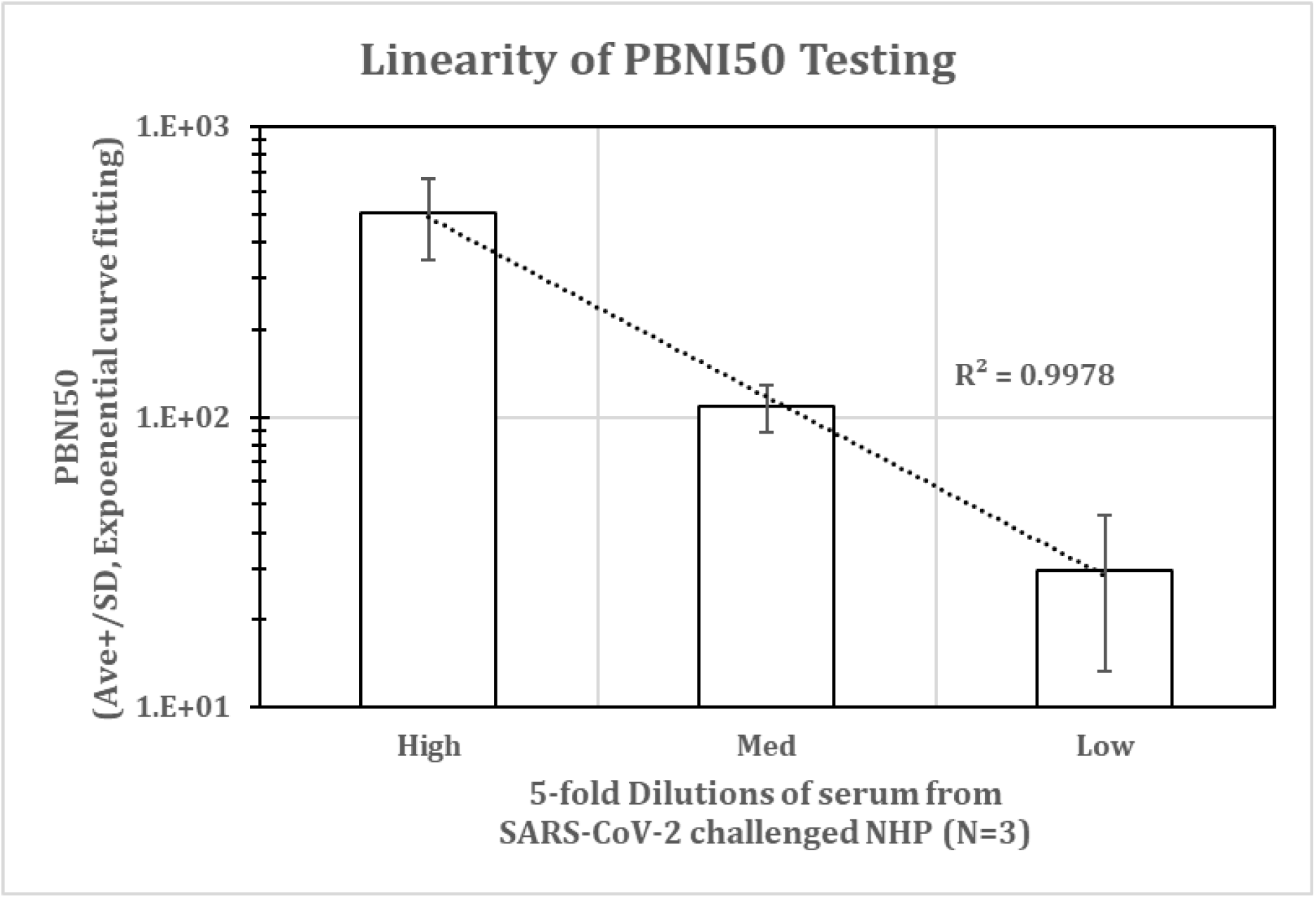
Linearity of PBNI_50_ testing: Three (3)-serum samples from SARS-CoV-2 challenged NHPs (1M16756, 3M16763, 3F16765) with high PBNI_50_ were serially diluted 5-fold to generate test samples with high, medium and low PBNI_50_, followed by PBNA with 6-point dilution in triplicates at each dilution. Average ± SD of the three PBNI_50_ for high, medium and low test samples are shown on the Y-axis. X-axis denotes the test samples with high, medium and low PBNI_50_. Trend analysis between the three groups is shown by the dotted line (R^2^=0.9978).

### 3.5. Comparison of different positive controls in PBNA to evaluate neutralization against variants of concern

A universal standard is necessary to harmonize the assays in different laboratories around the globe. Such a standard would facilitate the comparison of assay results from diverse laboratories. Currently, NIBSC 20/130 research reagent (anti-SARS-CoV-2 antibody) offered by NIBSC is intended to be used as a positive control for the development and evaluation of serological assays to detect antibodies against SARS-CoV-2. To evaluate the activity of the NIBSC 20/130 in PBNA, it was tested against all three pseudoviruses (614D, B1.357 RSA and B1.1.7 UK variant). Two other positive controls (Acro-SAD-S35 and Sino-40591-MM43) were also included in the assay. As shown in Figure. 5A and Table 1, NIBSC 20/130 demonstrated robust activity against 614D (PBNI_50_ of 3158.3±1,344.5), moderate activity against B1.1.7 UK variant (PBNI_50_ of 796.4 ± 127.5) and significantly poor activity against B1.351 RSA Variant (PBNI_50_ of <373.3 ± 369.5). Another positive control Acro-SAD-S35 (Figure.5B) showed robust activity against both 614D and B1.1.7 UK variant (PBNI_50_ of 10,764.5 ± 722.6 and 12,392 ± 4,577.1 respectively), but poor activity against B1.351 RSA variant (PBNI_50_ of <373.33 ± 369.5). In contrast, the monoclonal antibody Sino 40591-MM43 (Figure.5C) showed robust activity against all three variants; 614D, B1.351 RSA and B1.1.7 UK Variants (PBNI_50_ of 14,620.3 ± 946, 11.417.7 ± 2,912.1, 13,374.5 ± 5,118.8 respectively). These data suggest the differential neutralization of the positive controls (NIBS 20/130, Acro-SAD-S35 and Sino 40591-MM43) against 614D, B1.351 RSA and B1.1.7 UK Variants.

**Figure 5.**
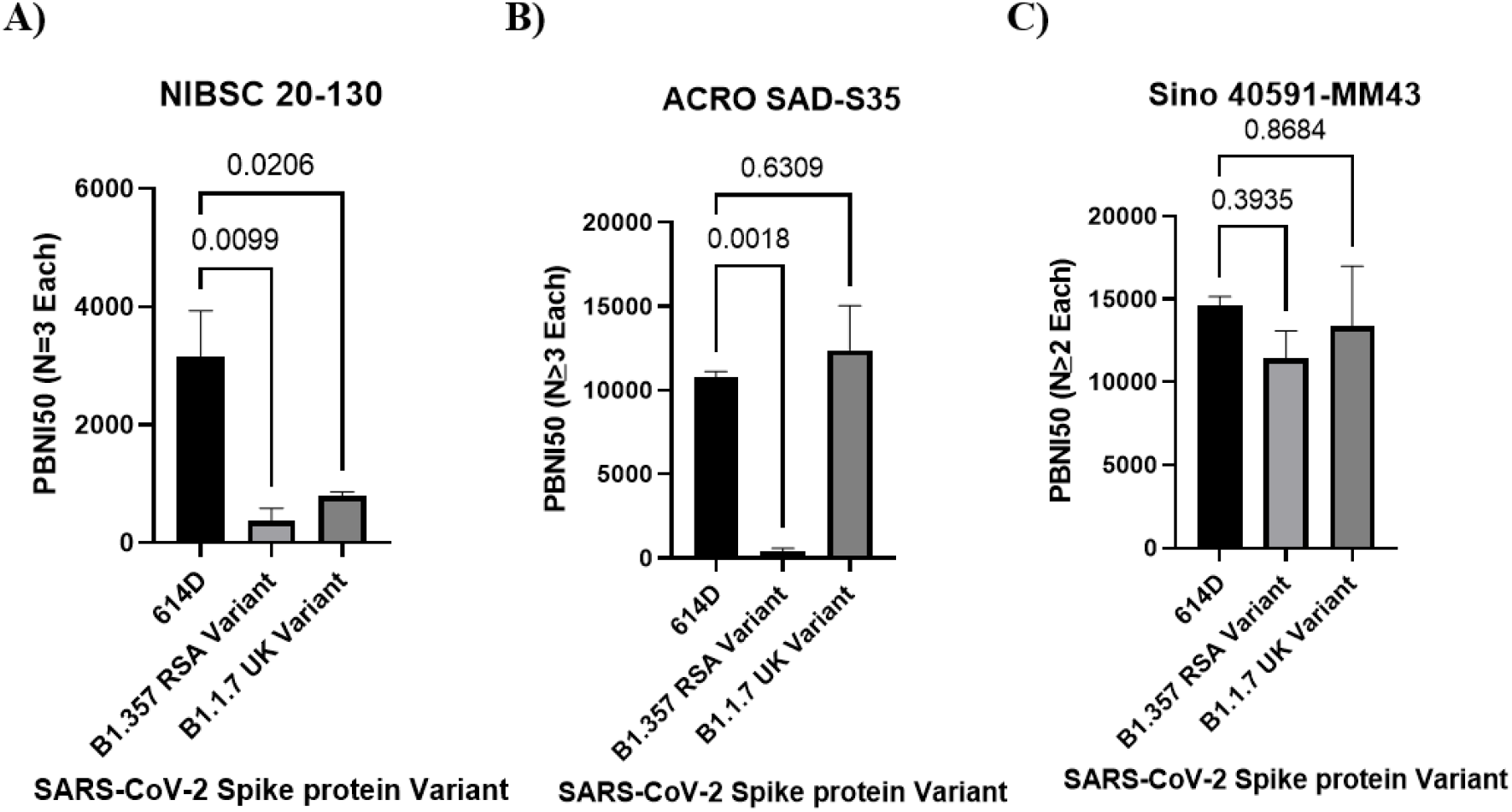
NIBSC 20/130 control shows lower neutralization against Variants of Concern (VOC) compared to 614D: Three positive controls (NIBSC 20/130, Acro SAD-S35, Sino 40591-MM43) were subjected to PBNA using the pseudo-viruses for 614D (A), B1.357 RSA Variant (B), B1.1.7 UK Variant (C) as described in the methods. Ave ± SD of PBNI_50_ from multiple experiments as indicated in the figure are shown on the Y-axis. Ordinary one-way analysis (multiple comparisons) was performed using GraphPad PRISM software. Adjusted p values are shown in the figure.

**Table 1.** PBNI_50_ data from independent experiments is shown in the table. A commercially available monoclonal antibody retains activity against SA and UK variants.

| Control | 614D | B.1.351 RSA Variant | B1.1.7 UK Variant |
| --- | --- | --- | --- |
| **NIBSC**<br>**20/130** | 4,270 | 160 | 667.1 |
|  | 3,541 | <800 | 800 |
|  | 1,664 | <160 | 922 |
| **Acro-**<br>**SAD-S35** | 11,511 | <160 | 11,192 |
|  | 10,502 | <800 | 8,535 |
|  | 11,164 | <160 | 17,450 |
|  | 9,881 |  |  |
| **Sino**<br>**40591-MM43** | 13,973 | 13,828 | 9,755 |
|  | 15,706 | 8,182 | 16,994 |
|  | 14,182 | 12,243 |  |

## 4. Discussion

Assay dynamic range helps in the appropriate differentiation of the vaccines with differing neutralization potencies. Besides measuring the 50% inhibition, a wide dynamic range also enables differentiation of robust inhibition (90%-99%) and can help rank order the vaccine candidates based upon their neutralization profiles. Dose dependent, two (2)-log dynamic range in the assay signal observed in our initial dose range testing experiments, for all three viruses along with the robust S/B ratio confirmed the suitability of this assay, though we consistently observed 1-log lower assay signal in cells infected with B1.351 RSA variant pseudovirus compared to either 614D and B1.351. It remains to be seen, if the lower signal in B1.351 RSA variant pseudovirus is either due to lower infection levels or if the assay conditions need to be further fine-tuned to increase the assay signal. However, as we observed sufficient 2-log dynamic range and S/B ratio needed for the confirmation of neutralization, we proceeded further with the evaluation of the test serum and positive controls.

As the goal of our research was to evaluate the suitability of PBNA for measuring the neutralization of preclinical samples, serum from 6-NHPs challenged with SARS-CoV-2 WAI strain were tested against 614D pseudovirus. Test serum samples and the positive controls used in the assay demonstrated a dose dependent inhibition of the assay signal (pseudovirus based luciferase signal) (data not shown), with a clear differentiation between the samples and positive controls. Interestingly, serum samples from 3-different species of monkeys (Rhesus macaques, Cynomolgus monkeys and African green monkeys) showed similar neutralizing index against 614D pseudovirus, albeit with differences between individual animals. As these results confirmed the suitability of PBNA, we further confirmed reproducibility of our experiments by systematically comparing the results from two independent experiments. A strong correlation (R^2^ of 0.96) suggested good reproducibility of PBNI_50_ generated in our PBNA. Such a strong assay reproducibility would facilitate the further validation of this assay for evaluating the clinical samples. Interestingly, we did not observe significant neutralization of B1.351 RSA variant with 5 out of 6 NHP challenged serum samples, despite having higher neutralizing index for both 614D and B1.1.7, UK variants. Our observation of similar neutralization of 614D and B1.1.7, UK variant, but not against B1.351 RSA variant by the majority of (5 out of 6 animals or 83.3% with lower neutralization against B1.351 RSA variant,) serum samples from SARS-CoV-2 challenged NHPs is not too surprising as a similar trend has been reported in the literature [29,30]. This similar trend between the preclinical (SARS-CoV-2 challenged NHP serum used in our experiments) and clinical samples (reported in the literature) increases the confidence in our assay system for its suitability to measure neutralizing antibodies against SARS-CoV-2.

It is also interesting to note that 5 out of 6 normal unexposed NHP samples tested (data not shown) in our assay showed high background neutralization index (PBNI_50_ of 100 to 250) against 614D. Only one of the NHPs showed a PBNI_50_ of <10. As we were not sure about the reason/specificity of this background PBNI_50_ level in the unexposed NHP serum samples, we serially diluted our SARS-CoV-2 challenged NHP serum samples with high PBNI_50_ levels (>500) to generate test samples with tiered PBNI_50_ levels (high, medium and low) followed by evaluation in PBNA. This experiment also enabled the measurement of linearity and lower limit of detection of neutralization index in PBNA. Our experiments confirmed the linearity of detection to approximately PBNI_50_ of 30 (the lowest PBNI_50_ tested in our assay), albeit with high %CV at the lower end of detection. As we confirmed the sensitivity of PBNA, high background levels (PBNI_50_ of 100 to 250) in 5 out of 6 unexposed NHP serum samples, might be due to pre-exposure to other coronaviruses in their natural environment (which might have generated neutralizing antibodies). Couple of previous reports about preexisting humoral immunity to SARS-CoV-2 in humans support this explanation [31,32].

In the literature, PBNA has been shown to correlate well with other assays measuring the neutralization antibodies; Plaque reduction neutralization test [33], microneutralization assay and ELISA assay [34]. Our results along with these previous reports provide a strong justification for the implementation of PBNA as a cost-effective measure of neutralizing antibodies in the preclinical studies of vaccine candidates.

Inclusion of a universally acceptable robust positive control is essential in the evolving serological/neutralization assays for SARS-CoV-2. Towards this goal, we used NIBSC 20/130 reagent as a positive control. However, in our experiments NIBSC 20/130 reagent showed impaired ability to neutralize the B1.1.7, UK and B1.351 RSA variants. In contrast, a monoclonal antibody (Sino 40591-MM43) tested as a positive control in our experiments retained robust activity against B1.1.7 and B1.351 RSA variants. This is one of the first reports showing the impaired ability of NIBSC 20/130 reagent to neutralize the B1.1.7, UK and B1.351 RSA variants. Demonstration of robust activity of a commercially available monoclonal antibody (Sino 40591-MM43) against 614D, B1.1.7 and B1.351 RSA variants is also a novel observation. It remains to be further confirmed, if the neutralization activity of Sino 40591-MM43 is retained against other variants of concern such as B1.617 delta and kappa variants. Nevertheless, as Sino 40591-MM43 is a commercially available monoclonal antibody, one could envision accessibility across the globe with unlimited supply (as it can be scaled up due to the monoclonal nature). This could further enhance the global harmonization of the neutralization and serological assays against SARS-CoV-2 variants of concerns.

## 5. Conclusions

Our findings demonstrate successful development of a robust pseudovirus based neutralization assay against SARS-CoV-2 variants of concern, which can aid evaluation of preclinical and clinical samples and rapidly develop effective countermeasures against SARS-CoV-2 and the variants of concern. Our results also identified a commercially available monoclonal antibody that can serve as a potential global standard in the neutralization assays against SARS-CoV-2 variants of concern.

## Author Contributions

Conceptualization, R.K. and F.K.; methodology, Z.C., S.L. and Y.V.K.; writing—original draft preparation, R.K., Y.V.K. and Z.C.; writing—review and editing, R.K., F.K. and J.F. All authors have read and agreed to the published version of the manuscript.

## Ethical Guidelines for the Use of Animals in Research

All procedures involving animals were approved by Southern Research Institutional Animal Care and Use Committee (IACUC). This study was carried out in strict accordance with the recommendations in the Guide for the Care and Use of Laboratory Animals [35]

## Acknowledgments

The opinions or assertions contained herein are the private views of the authors and are not necessarily those of Southern Research. The following are acknowledged for their support: National Institute of Allergy and Infectious Diseases (NIAID) for the permission to use the serum samples from the project “infectivity study of sars-cov-2 in rhesus macaques (RM), Cynomolgus macaques (CM), and African green monkeys (AGMs) challenged via single respiratory route of infection” (SR project 15891.08.01). Jennifer Pickens for sharing the serum samples from the SARS-CoV-2 challenged NHP. NHP serum used in the experiment was generated under National Institutes of Health (NIH)/National Institute of Allergy and Infectious Diseases (NIAID) contract HHSN272201700029I; B08 Task Order No.75N93020F00001; Development and Use of a Non-Human Primate Model of SARS-CoV-2 Infection.

